# Virucidal Star Polymer Disrupts Alphavirus Infection in Aedes Mosquitoes

**DOI:** 10.64898/2026.11.11.765866

**Authors:** Shannan-Leigh Macleod, Maeva Van Langendonck, Noémie Rase, Ana L. Rosales-Rosas, Samuel T. Jones, Leen Delang

**Affiliations:** KU Leuven, Department of Microbiology, Immunology and Transplantation, Rega Institute for Medical Research, VITA, 3000 Leuven, Belgium; Department of Materials and Henry Royce Institute, University of Manchester, Oxford Road, Manchester, M13 9PL, United Kingdom; School of Chemistry, University of Birmingham, Edgbaston, Birmingham, B15 2TT, United Kingdom

## Abstract

Mosquito-borne alphaviruses represent an increasing global health threat, yet there are currently no antiviral interventions capable of targeting both the vertebrate host and mosquito vector. Here, we investigated the antiviral potential of the heparan sulfate proteoglycan-mimicking zwitterionic polymer ZP12 against alphavirus infection in *Aedes aegypti* mosquitoes. ZP12 exhibited potent antiviral and virucidal activity against Ross River virus (RRV) and Semliki Forest virus (SFV) in C6/36 mosquito cells. Antiviral activity was retained in *ex vivo* mosquito gut cultures, where both high (100 µg/mL) and low (0.5 µg/mL) concentrations significantly reduced RRV replication, with viral titres reduced by up to five orders of magnitude at peak infection. Delayed-treatment studies demonstrated that ZP12 retained antiviral activity after infection had been initiated, although efficacy declined as treatment was delayed. To evaluate prophylactic delivery, mosquitoes were exposed to ZP12 through sugar feeding prior to viral challenge. While this approach did not directly protect against subsequent gut infection, follow-up experiments demonstrated that ZP12 remained biologically active within the mosquito, suggesting that limited delivery to the midgut rather than loss of antiviral activity restricted efficacy. Collectively, these findings demonstrate that ZP12 is a potent antiviral and virucidal agent against alphaviruses in mosquito cells and tissues and supports its further development as a transmission-blocking strategy for arboviral control.

## Introduction

Arboviruses are a diverse group of viruses primarily transmitted to vertebrate hosts by arthropod vectors such as mosquitoes. Among these, dengue virus and chikungunya virus (CHIKV) are of significant global concern. Alarmingly, the incidence of infections caused by these arboviruses is rising and increasingly being recognised as a potential driver of the next global pandemic^1^. This growing threat is largely attributed to the expanding range of competent vectors, a trend driven by changes in climate, globalisation and urbanisation^2–5^.

Arboviral transmission is primarily mediated by female *Aedes (Ae.) aegypti* mosquitoes^6^. Following ingestion of an infectious blood meal, viruses establish infection in the mosquito midgut before disseminating to secondary tissues including the salivary glands, enabling subsequent transmission to a new host^7,8^. Because the mosquito midgut is the initial site of viral replication, *ex vivo* gut cultures provide a physiologically relevant platform for studying virus-vector interactions and evaluating antiviral interventions while retaining native tissue architecture^9,10^.

Heparan sulphate proteoglycans (HSPGs) serve as common attachment factors for a wide range of viruses, including mosquito-borne alphaviruses, during host cell attachment and entry. This virus-receptor interaction has become a focal point for the development of novel therapeutic approaches. Recently, a new class of extracellular antivirals has emerged based on HSPG mimetics, including star-shaped antiviral polymers (star-polymers). These compounds competitively inhibit viral attachment and exert virucidal activity through multivalent interactions with viral surface proteins^11–15^. Importantly, mosquito-borne alphaviruses, including CHIKV, rely on HSPG receptors within mosquito midgut and salivary gland epithelial cells during early stages of infection^16^. This shared reliance on HSPGs in both vertebrate and mosquito hosts raises the possibility of a dual-action antiviral capable of limiting infection in both settings.

We recently developed the star zwitterionic polymer ZP12^17^, which exhibits broad-spectrum virucidal activity against multiple viral families, including *Alphaviridae, Flaviviridae, Herpesviridae and Picornaviridae*. Furthermore, ZP12 demonstrated potent therapeutic effects in *in vivo* murine models of CHIKV infection^17^. ZP12 significantly reduced joint swelling, decreased tissue viral load by 90%, and attenuated activation of immunopathogenic pathways associated with CHIKV-induced inflammation. To determine whether ZP12 could also interfere with viral replication in the mosquito vector, we next evaluated ZP12 in the *ex vivo* mosquito gut model^9^. Using this model, we investigated the antiviral activity of ZP12 against Ross River virus (RRV), a closely related alphavirus to CHIKV, to assess its ability to limit viral replication within mosquito gut tissues.

## Results and Discussion

### Virucidal Properties of ZP12 in Mosquito Cells

In mammalian cell lines (Vero and HEK293T), ZP12 demonstrated low toxicity and potent antiviral activity against CHIKV and ONNV, with an IC_50_ of approximately 0.55 *µ*M and a selectivity index (SI) of ~13^17^. As no detectable cytotoxicity was observed across the concentration range tested, SI values (CC_50_/IC_50_) were conservatively estimated using 300 μg/mL as the CC_50_ threshold, consistent with previous antiviral polymer studies^11,12,18–22^. Given that both mammalian and mosquito cells utilise HSPGs during viral attachment, we next investigated the biocompatibility and antiviral activity of ZP12 in C6/36 mosquito cells. MTS analysis confirmed that ZP12 was non-toxic, even at the highest concentration tested (5000 *µ*g/mL; SI Fig. 1).

RRV and Semliki Forest virus (SFV) were selected representative alphaviruses due to their close genetic and structural similarity to CHIKV, including their reliance on HSPGs during host cell entry^23^. To evaluate antiviral activity, ZP12 was pre-incubated with either RRV or SFV prior to inoculation onto C6/36 cell monolayers, and infectivity was quantified by flow cytometry at 72 hours post infection (hpi) (Fig. 1). ZP12 reduced infection by both viruses in a dose-dependent manner. Notably, ZP12 exhibited greater antiviral potency in mosquito cells than previously observed in mammalian cells, with IC_50_ values of approximately 0.34 μM (SI 21) and 1.47 μM (SI 204) were obtained for RRV and SFV, respectively (Fig. 1a-b). The enhanced activity observed in mosquito cells may reflect differences in HSPG dependence and viral entry mechanisms between mosquito and mammalian hosts. Although differences in potency were observed between RRV and SFV, potentially reflecting varying degrees of HSPG utilisation, ZP12 demonstrated potent antiviral activity against both viruses.

**Fig. 1:**
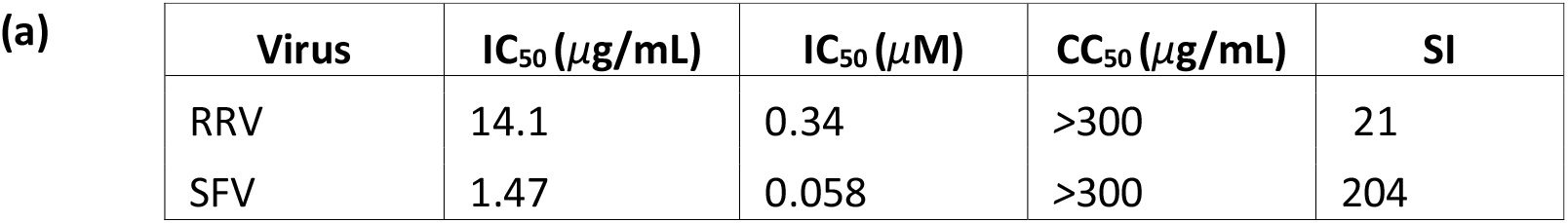
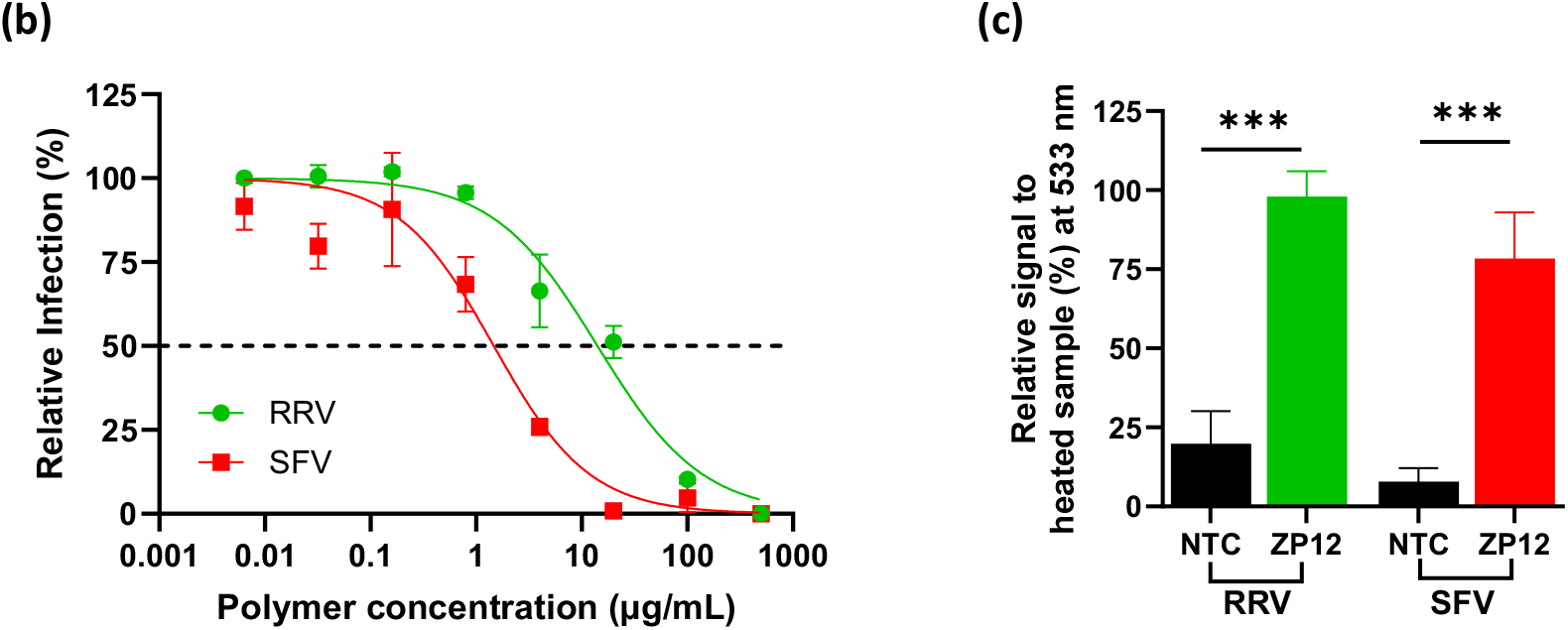
Virucidal antiviral activity of ZP12 against RRV and SFV in C6/36 mosquito cells. **(a)** Summary of antiviral activities. **(b)** Dose response curves showing inhibition of RRV and SFV infection following ZP12 treatment. Percentage infection was normalised to untreated virus controls, with the dashed line indicating 50% inhibition. **(c)** Assessment of virucidal activity using FAIRY. Fluorescence signals were normalised to heat-treated virus controls, representing 100% genome exposure. Increased fluorescence indicates viral RNA release following virion disruption. Data are presented as means±SEM from at least duplicate samples and are representative of three independent experiments (n = 3). Comparison between groups were performed with non-parametric Mann-Whitney U test (***p<0.001). Key: IC_50,_ 50% inhibitory concentration; CC_50,_50% cytotoxicity concentration; SI, selectivity index (CC_50_/IC_50_); NTC, non-treated control.

To determine whether ZP12 possesses a virucidal mode of action, viral integrity was assessed using the Fluorescence Assay for vIRal IntegritY (FAIRY)^24^. This cell-free assay quantifies viral RNA release from damaged virions through binding of a fluorescent nucleic acid probe to exposed RNA genomes. An increase in fluorescence therefore reflects loss of virion integrity and provides an indirect measure of viral inactivation. Following ZP12 treatment, fluorescence increased by approximately 74% for both RRV and SFV, indicating substantial viral RNA release consistent with virion disruption and a virucidal mechanism of action (Fig. 1c). These findings are consistent with previous reports demonstrating the ability of sulfonated star-polymers to irreversibly inactivate enveloped viruses through direct virion damage^11,13,14^. Collectively, these findings demonstrate that ZP12 exhibits potent antiviral and virucidal activity against alphaviruses in mosquito cells, supporting its potential for limiting arboviral replication within the mosquito vector.

### Viability of ZP12-treated *Ex Vivo* Gut Cultures

Following the demonstration of potent antiviral activity and low toxicity in mosquito cell lines, we next evaluated the suitability of ZP12 for use in established *ex vivo* mosquito gut cultures, a physiologically relevant model of alphavirus infection within the mosquito vector^25^. Prior to antiviral testing, ZP12 was evaluated at concentrations up to 100 µg/mL to ensure tissue saturation and identify any potential toxic effects under conditions exceeding those expected to be required for antiviral activity.

To assess tissue viability, dissected Ae. aegypti mosquito guts were cultured in the presence of ZP12 for three days. Spontaneous hindgut contractions were qualitatively observed throughout the culture period, suggesting that tissue viability had been maintained following dissection. Quantitative assessment using the CellTiter-Glo 3D ATP assay further confirmed that metabolic activity was maintained across all treatment groups, including the highest concentration tested (100 µg/mL) (Fig. 2a). These findings demonstrate that ZP12 is well tolerated by *ex vivo* mosquito gut tissues, with no detectable toxicity observed at concentrations up to 100 µg/mL. Having established the biocompatibility of ZP12, both a high concentration (100 µg/mL) and a lower concentration (0.5 µg/mL) were subsequently evaluated for antiviral activity in infected gut cultures.

**Fig. 2:**
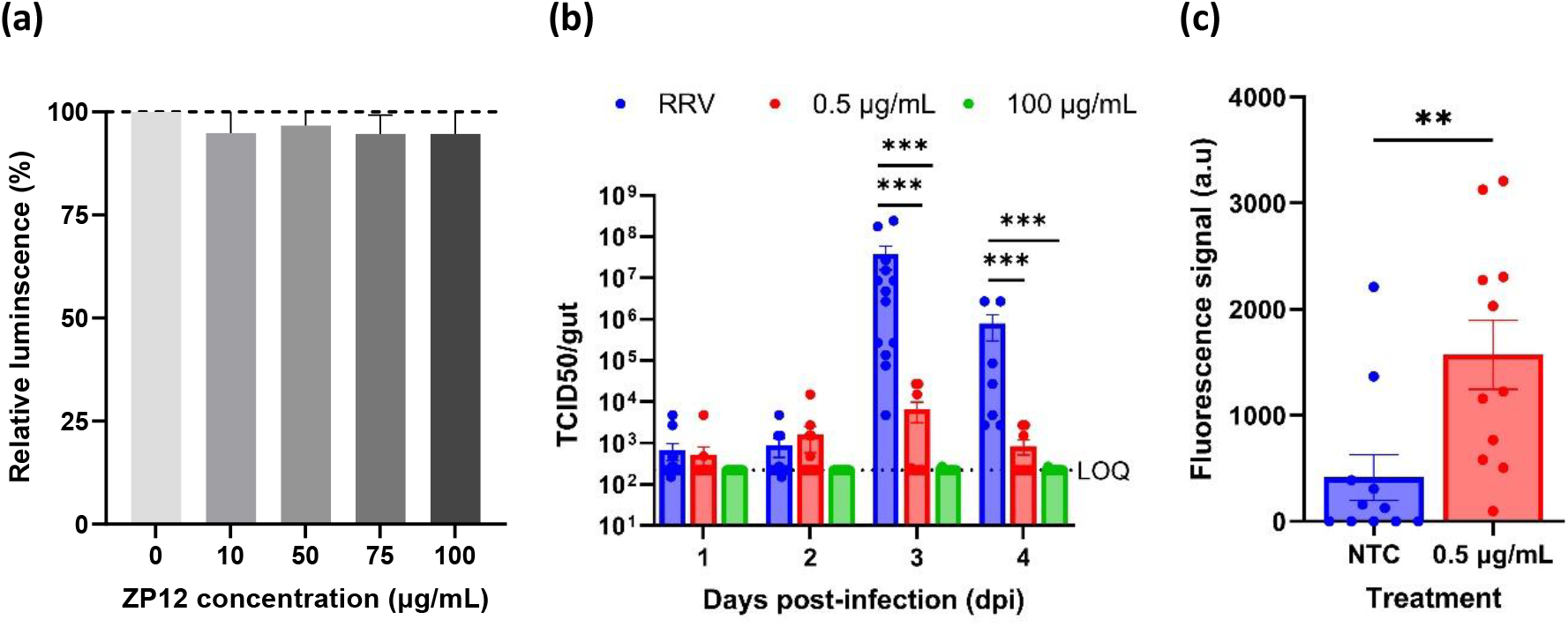
Antiviral activity of ZP12 in ex vivo RRV-infected gut cultures. **(a)** Viability of mosquito gut tissues following 3 days of exposure to ZP12. Viability was normalised to untreated control guts. **(b)** Infectious viral replication kinetics in RRV-infected gut cultures treated with low (0.5 µg/mL) or high (100 µg/mL) concentrations of ZP12, as determined by end-point titration. The dashed line indicates the limit of quantification (LOQ). **(c)** Assessment of virucidal activity following treatment with 0.5 µg/mL ZP12 by FAIRY at 3 dpi. Increased fluorescence indicates viral genome exposure following virion disruption. Data are presented as mean±SEM from 4-6 mosquito guts per group and are representative of at least two independent experiments (n=2). Each dot represents an individual mosquito gut. Comparison between groups were performed with non-parametric Mann-Whitney U test (two-tailed **p<0.01 and ***p<0.001).

### Antiviral Activity of ZP12 in RRV-infected *Ex Vivo* Mosquito Gut Cultures

Next, the antiviral activity of ZP12 was evaluated in RRV-infected *ex vivo* gut cultures. RRV was selected as a surrogate alphavirus model due to its close relationship to CHIKV and the availability of a well-established *Ae. aegypti ex vivo* gut infection model^25^. Infected gut cultures (10^5^ PFU/mL) were treated with either a low concentration (0.5 µg/mL) or high concentration (100 µg/mL) of ZP12 and monitored for up to 4 dpi by end-point titration. In untreated cultures, RRV replicated efficiently, reaching peak titres at 3 dpi before declining, consistent with previously reported replication kinetics^25^ (Fig. 2b). ZP12 treatment reduced viral replication in the low (0.5 µg/mL) and high (100 µg/mL) concentrations by approximately 3.7 logs and 5 logs, respectively, at 3 dpi relative to untreated controls. These findings demonstrate potent antiviral activity across a broad concentration range and indicate that substantial inhibition of viral replication can be achieved even at low polymer concentrations. As future studies will evaluate ZP12 in live mosquitoes, where total polymer exposure is limited by the small volume of sugar meal consumed, subsequent analyses will focus on the lower concentration (0.5 µg/mL). Despite being 200-fold lower than the maximally effective dose, this concentration retained strong antiviral activity, highlighting its potential translational relevance for *in vivo* mosquito applications.

To determine whether this antiviral effect was associated with virion disruption, viral integrity was assessed using FAIRY^24^. Following treatment with 0.5 ug/mL, a significant increase in fluorescence was observed relative to untreated controls (Fig. 2c), indicating increased viral genome exposure consistent with disruption of virion integrity. These findings demonstrate that the antiviral activity observed at biologically relevant concentrations is associated with a virucidal mechanism of action. Collectively, these findings demonstrate that ZP12 potently suppresses RRV infection in *ex vivo* mosquito gut cultures. The marked reduction in infectious virus production, together with increased viral genome exposure detected by FAIRY, provides complementary evidence that ZP12 inhibits alphavirus infection within mosquito tissues through a virucidal mechanism of action.

### Delayed Treatment of *Ex Vivo* Cultured Guts

To investigate whether ZP12 could suppress ongoing viral replication, the *ex vivo* mosquito gut model was adapted to evaluate delayed treatment regimens. Whereas previous experiments assessed antiviral activity immediately following viral inoculation, here ZP12 treatment was initiated at different stages of infection to determine the therapeutic window of antiviral activity (Fig. 3a). RRV-infected gut cultures (10^5^ PFU/mL) were treated with ZP12 at either 1, 2 or 3 dpi, and infectious viral titres were quantified 24 hours later by end-point titration. For the 1 dpi treatment group, an additional collection was performed at 48 hours post-treatment (3 dpi) to assess the effect of prolonged exposure.

**Fig. 3:**
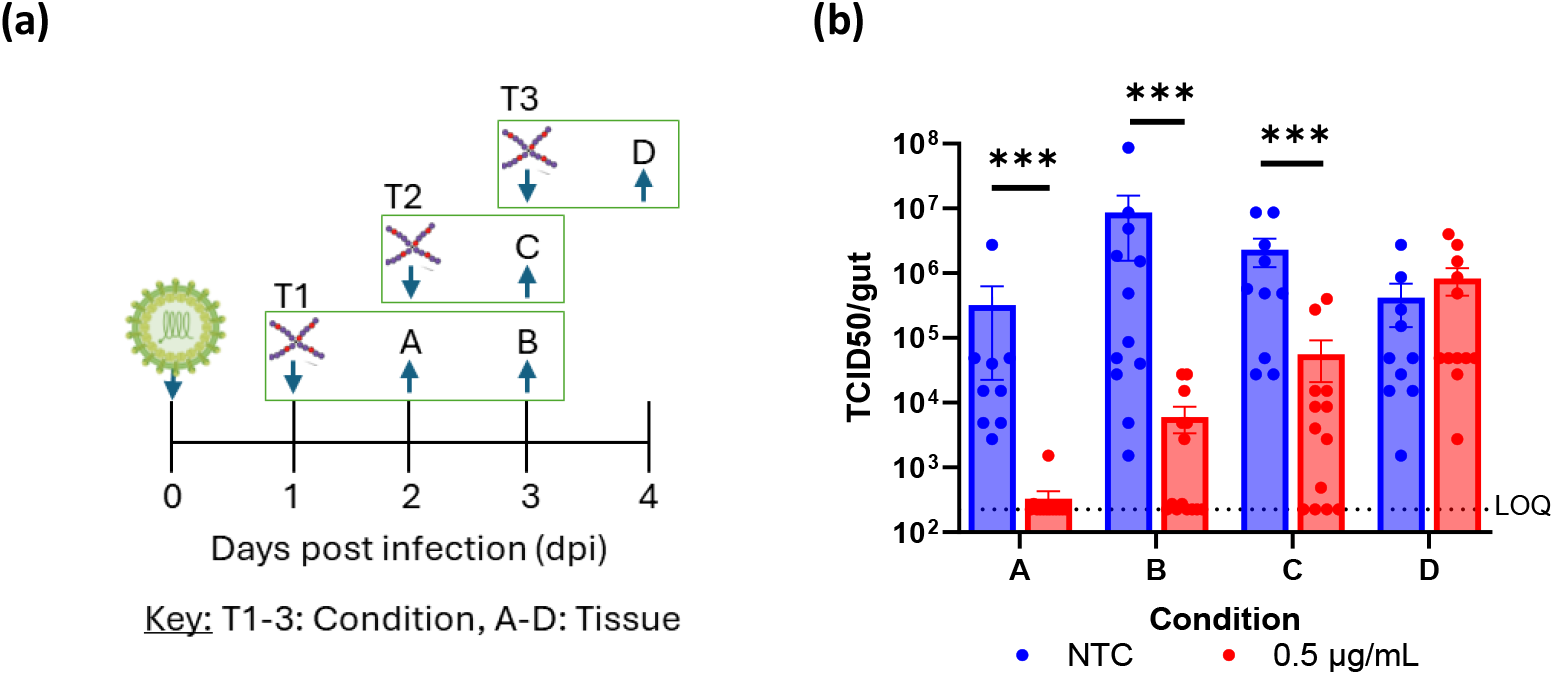
Effect of ZP12 after established RRV infection in ex vivo cultured guts. **(a)** Schematic overview of delayed-treatment conditions (T1-T3) and corresponding gut collection time points (A-D). **(b)** Infectious viral titres in RRV-infected gut cultures treated with ZP12 at different stages of infection, compared with untreated controls. The dashed line indicates the limit of quantification (LOQ). Data are presented as mean ± SEM from 4-6 mosquito guts per experimental group and are representative of three independent experiments (n = 3). Each dot represents an individual mosquito gut. Statistical comparisons between groups were performed using a Mann-Whitney U test (two-tailed ***p<0.001).

Treatment initiated at 1 dpi (Condition A) significantly reduced viral titres by approximately 2.5 logs within 24 hours (2 dpi), indicating that ZP12 retained antiviral activity after viral replication had already been initiated within the tissue. Extending treatment for a further 24 hours enhanced this effect (Condition B), resulting in an approximately 3-log reduction in infectious titre at 3 dpi (Fig. 3b). When treatment was delayed until 2 dpi (Condition C), viral titres were still reduced by approximately 1.6 logs compared with untreated controls, although the magnitude of inhibition was lower than that observed following treatment at 1 dpi (Condition A and B). In contrast, treatment initiated at 3 dpi (Condition D) had no significant effect on viral titres. Together, these findings indicate that antiviral efficacy progressively declines as infection advances, with the greatest reduction in viral burden achieved when treatment is administered before peak viral replication.

The reduced efficacy observed following delayed treatment is consistent with the proposed virucidal mechanism of ZP12. As a direct-acting extracellular antiviral, ZP12 primarily targets free virions and prevents subsequent rounds of infection. Consequently, its ability to limit viral spread is expected to diminish as infection progresses and the number of infected cells increases.

Collectively, these findings demonstrate that ZP12 retains antiviral activity when administered after viral replication has been initiated, although efficacy is strongly dependent on treatment timing. These findings support the potential use of ZP12 as a prophylactic or early transmission-blocking strategy, for example through sugar-feeding formulations encountered shortly after acquisition of an infectious blood meal^26,27^. Early exposure to ZP12 could suppress viral amplification within the mosquito midgut and thereby reduce the likelihood of subsequent virus transmission.

### Prophylactic Treatment of *Ae. aegypti* Mosquitoes

Given that ZP12 exhibited the greatest antiviral efficacy when administered during the early stages of infection, we next investigated whether prophylactic exposure could protect mosquitoes against subsequent viral challenge. To mimic a potential field-based delivery strategy, adult *Ae. aegypti* mosquitoes were exposed to ZP12 through a sugar meal for 24 h prior to gut dissection. This approach was designed to assess whether pre-existing levels of ZP12 within the mosquito could suppress viral replication following exposure to RRV.

Unlike the *ex vivo* gut culture model, where gut tissues can be directly exposed to defined concentrations of ZP12, oral delivery in live mosquitoes is constrained by the limited volume of sugar meal consumed (~5 µL per mosquito per day)^28^. Consequently, mosquitoes were provided with a ZP12-containing sugar solution (100 ng/µL), corresponding to an estimated intake of approximately 0.5 µg per mosquito. This exposure was selected because substantial antiviral activity had already been observed at low ZP12 concentrations (*ex vivo*), suggesting that biologically relevant antiviral effects may still be achievable following oral administration.

Following this pre-feeding prophylactic exposure period, mosquito guts were dissected and infected *ex vivo* with RRV (1×10^5^ PFU/mL) to determine whether prior ingestion of ZP12 conferred antiviral protection. Unexpectedly, no significant differences in viral titres were observed between ZP12-treated and untreated mosquitoes gut cultures at either 1, 2 or 3 dpi (Fig. 4). These findings suggest that prior ingestion of ZP12 did not confer measurable antiviral protection within the gut tissue.

**Fig. 4:**
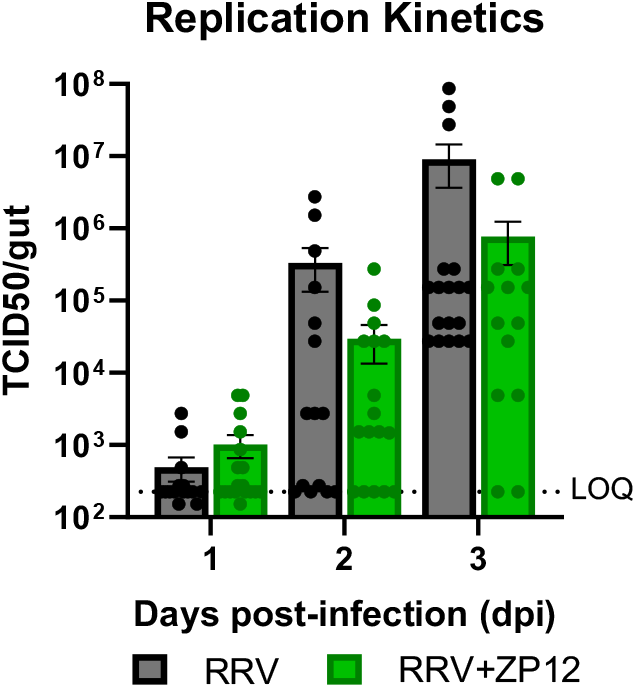
Evaluation of prophylactic ZP12 treatment in Ae. aegypti mosquitoes. Adult mosquitoes were fed a ZP12-containing sugar meal for 24 h prior to gut dissection and subsequent ex vivo RRV infection. Infectious viral titres were measured at the indicated time points by end-point titrations. The dashed line indicates the limit of quantification (LOQ). Data are presented as mean±SEM from 4-6 mosquito guts per experimental group and are representative of two independent experiments (n=2). Each dot represents an individual mosquito gut. Statistical comparisons were performed using a two-tailed Mann-Whitney U test.

To determine whether the lack of efficacy resulted from loss of antiviral activity following ingestion, whole mosquitoes previously exposed to ZP12 were homogenised and the resulting lysates were overlaid onto untreated RRV-infected *ex vivo* gut cultures (Fig. 5a). Viral titres were assessed at 3 dpi. Notably, treatment with whole-mosquito lysates resulted in an approximately 1.6-log reduction in viral titre compared with untreated controls, demonstrating that ZP12 remained biologically active within the mosquito following oral administration. These findings suggested that the lack of prophylactic efficacy was not due to degradation or excretion of ZP12 and warranted further investigation into its distribution within the mosquito.

**Fig. 5:**
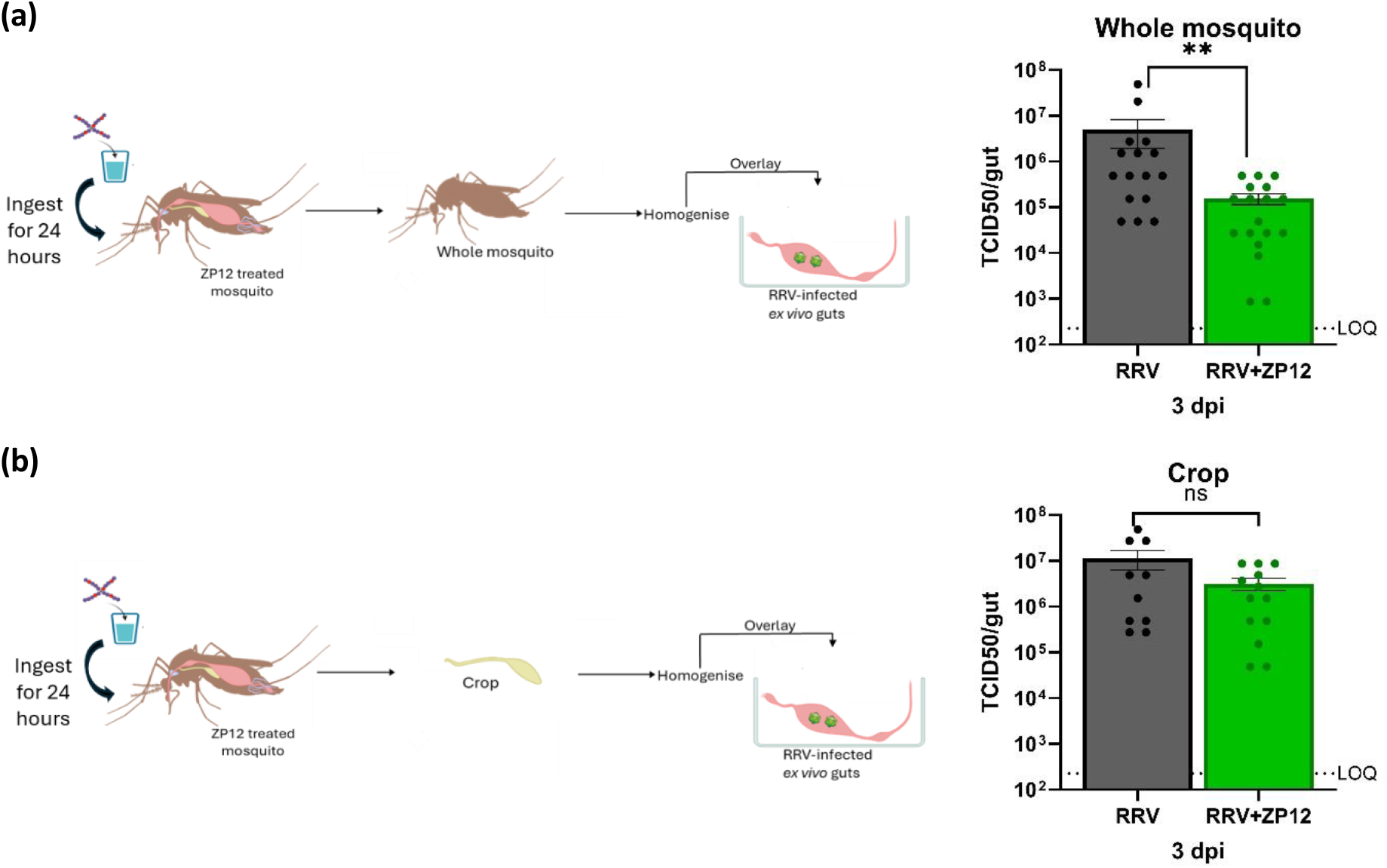
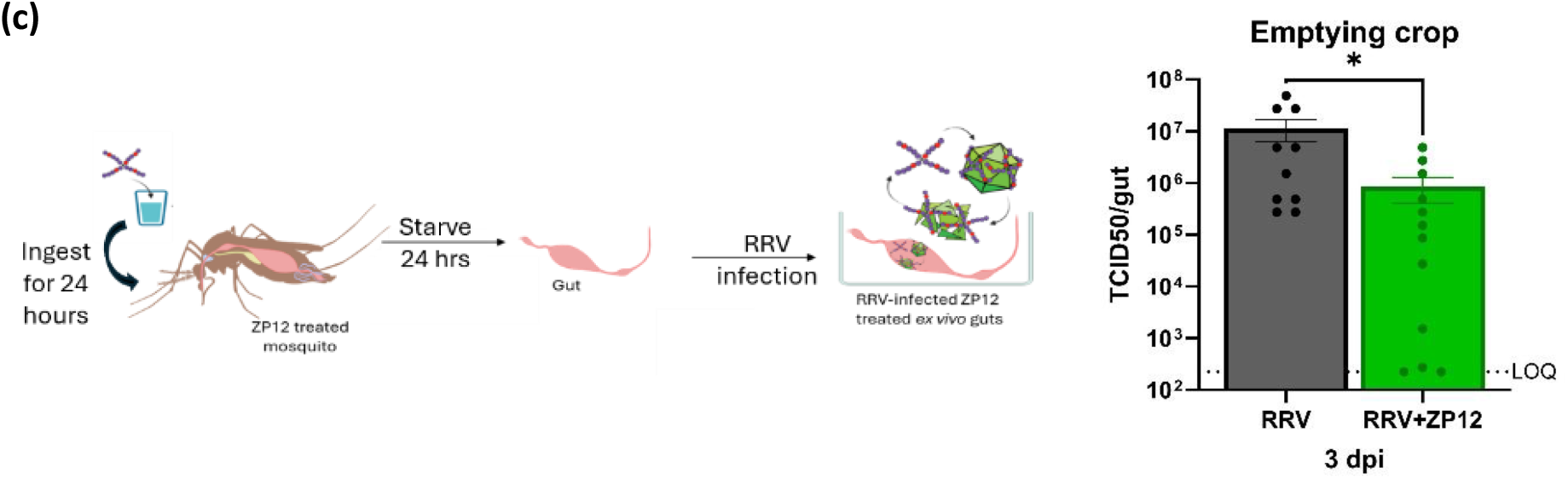
Evaluation of prophylactic ZP12 treatment in Ae. aegypti mosquitoes. **(a)** Whole-mosquito lysates from ZP12-fed mosquitoes were overlaid onto RRV-infected ex vivo gut cultures. **(b)** Crop lysates from ZP12-fed mosquitoes were overlaid onto RRV-infected ex vivo gut cultures. **(c)** Mosquitoes were starved following ZP12 feeding prior to gut dissection and ex vivo RRV infection. Infectious viral titres were quantified at 3 dpi. Dashed line indicates the limit of quantification (LOQ). Data are presented as mean±SEM and are representative of two independent experiments (n=2). Each dot represents an individual mosquito gut. Statistical comparisons were performed using a Mann-Whitney U test (two-tailed *p<0.05, **p<0.01).

In mosquitoes, sugar meals are primarily stored within the crop, whereas blood meals are delivered directly to the midgut, the primary site of alphavirus replication^29^. We therefore hypothesised that ZP12 was being retained within the crop and was unable to reach the gut at sufficient concentrations to exert antiviral activity (Fig. 5b). To test this hypothesis, crops from ZP12-fed mosquitoes were dissected, homogenised and overlaid onto untreated RRV-infected *ex vivo* gut cultures. Viral titres were again assessed at 3 dpi. However, no significant differences were observed between crop lysate-treated and untreated cultures. Given the technical difficulty associated with crop dissections, this result may have been influenced by leakage or rupture of the crop during isolation, resulting in loss of crop contents prior to analysis.

To further investigate whether sequestration within the crop was limiting gut exposure, mosquitoes were starved following ZP12 feeding to promote mobilisation of stored sugar reserves and facilitate transfer of crop contents to the gut (Fig. 5c). Under these conditions, an approximately 1-log reduction in viral titre was observed compared with untreated controls, suggesting that redistribution of ZP12 from the crop may partially restore antiviral activity within the gut. Collectively, these findings indicate that the lack of prophylactic efficacy observed following sugar feeding was not due to loss of ZP12 antiviral activity within the mosquito. Instead, the data suggest that sequestration of ZP12 within the crop limits its availability at the midgut, the primary site of viral replication. The recovery of antiviral activity following mosquito starvation further supports this interpretation and highlights tissue distribution, rather than antiviral potency, as the principal barrier to effective oral delivery. Future studies should therefore focus on optimising delivery strategies that enhance transfer of ZP12 from the crop to the midgut, thereby improving its potential as a prophylactic transmission-blocking antiviral.

## Conclusion

Arboviruses, particularly mosquito-borne alphaviruses, continue to pose an increasing global health threat, highlighting the need for antiviral strategies that target both vertebrate hosts and mosquito vectors. Given the importance of HSPGs in alphavirus entry, HSPG-mimicking compounds such as ZP12 represent a promising broad-spectrum antiviral approach.

ZP12 exhibited potent antiviral and virucidal activity against the alphaviruses RRV and SFV in C6/36 mosquito cells, with greater potency than previously observed in mammalian systems. This activity was retained in *ex vivo* mosquito gut cultures, where both high and low concentrations significantly reduced viral replication, resulting in titre reductions of up to five orders of magnitude and evidence of virion disruption. Notably, ZP12 remained effective when administered after infection had been established, although antiviral efficacy declined as infection progressed. To investigate its potential as a prophylactic intervention, ZP12 was delivered to mosquitoes through sugar feeding prior to viral challenge. Although this approach did not reduce subsequent viral replication in infected gut tissues, the recovery of antiviral activity from whole-mosquito lysates and its partial restoration following starvation demonstrated that the compound remained biologically active within the mosquito. Together, these findings suggest that sequestration within the crop, rather than loss of potency, limits oral efficacy and identify tissue distribution as a key challenge for future delivery strategies.

Collectively, these findings establish ZP12 as a potent antiviral against alphaviruses in mosquito systems and identify tissue distribution as the main challenge for prophylactic application. Future studies should evaluate ZP12 in live mosquito infection models and develop delivery strategies that overcome crop sequestration, supporting its potential as a transmission-blocking intervention for arboviral disease control.

## Methodology

### Materials and general methods

Reagents were purchased from Gibco (ThermoFisher Scientific), unless otherwise stated. Biological study reagents were handled using aseptic techniques in class II BSC. ZP12 was dissolved in PBS and sterilised using a 0.22 *µ*m filter. Tecan Spark Multimode Microplate reader (Spark Control Magellan, v2.2) was used for all absorbance and luminescence readings.

### In Vitro Studies

#### 1. Cell culture

African green monkey kidney Vero cells (CCL-81, ATCC) were propagated in Minimum Essential Medium (MEM) supplemented with 10% heat-inactivated foetal bovine serum (FBS; HyClone), 1% L-glutamine, 1% sodium bicarbonate, 1% non-essential amino acids (NEAA) and 1% Penicillin-Streptomycin (P/S). C6/36 cells (CRL-1660, ATCC), were propagated in Leibovitz’s L-15 medium (Gibco), supplemented with 10% FBS, 1% NEAA, 1% HEPES buffer (Gibco) and 1% P/S. Cells were mycoplasma free and maintained at 37°C with 5% CO_2_ (Vero cells), and 28°C in a CO_2_ free environment (C6/36 cells). Virus propagation, antiviral assays and end-point titrations were performed using the media defined above but supplemented with 2% instead of 10% FBS, for both mammalian and mosquito cells.

#### 2. Viruses

RRV was obtained from the National Collection of Pathogenic Viruses (UK; catalogue number 0005281v). SFV (Vietnam strain; GenBank EU350586.1) belongs to the collection of the Rega Institute of Medical Research, Belgium. RRV-GFP reporter virus and SFV-mCherry was kindly provided by Prof. Mark T. Heise (University of North Carolina) and Prof. Andres Merits (University of Tartu, Estonia), respectively.

#### 3. C6/36 cell viability

Cell viability was assessed using CellTiter 96® AQueous One Solution (MTS) assay (Promega). C6/36 cells were seeded (1×10⁶ cells/well) and incubated overnight at 28°C. Cells were treated in triplicate with serial dilutions of ZP12 (50-5000 µg/mL). Untreated wells containing cells and medium served as negative controls, whilst blank wells containing medium only served as background controls. After 24 h, cell viability was assessed according to the manufacturer’s instructions. Briefly, medium was replaced with 100 µL MTS reagent and incubated for 1 h at 28°C in the dark. Absorbance was measured at 498 nm using a Tecan Spark plate reader. CC_50_ values were calculated using GraphPad Prism.

#### 4. Viral inhibition assay

C6/36 cells were seeded (1×10^6^ cells/well) in 24-well plates overnight. Serial dilutions of ZP12 (51.2 pg/mL-500 µg/mL) were pre-incubated with RRV-GFP or SFV-mCherry at an MOI of 1 for 1 h at 28°C prior to inoculation onto cell monolayers. Following a 1 h adsorption period, the inoculum was removed and replaced with fresh medium. Cells were incubated for a further 72 h before analysis by flow cytometry. Cells were stained with Zombie Aqua Fixable Viability Dye (Invitrogen), fixed with 4% paraformaldehyde (PFA), and acquired using a BD Fortessa (BD FACSDiva v8.0.2). GFP and mCherry expression were used as measures of infection. IC_50_ values were calculated by normalising infection levels to untreated virus controls and fitting dose-response curves using GraphPad Prism.

#### 5. FAIRY assay

FAIRY serves as a nucleic acid fluorescent readout for viral infectivity and is used to assess virion integrity.^277–279,402^ Briefly, ZP12 and RRV or SFV were mixed at a 1:1 ratio in phenol red-free DMEM without FBS and incubated for 1 h at 37°C. PBS-treated virus served as the negative control, whilst virus heated at 90°C for 10 min served as the positive control. Following incubation, equal volumes (50 µL) of sample and TO-PEG were combined in black flat-bottom 96-well plates and incubated for 10 min at room temperature. Fluorescence was measured using a Tecan Spark plate reader (*λ*_Ex_=510 nm and *λ*_Em_=533 nm). Fluorescence values were normalised to the heat-treated control, which was assumed to represent complete genome exposure.

### *Ex vivo* Gut Culture Studies

#### 6. Ae. aegypti rearing

Eggs belonging to the *Ae. aegypti* Paea strain (Papeete, Tahiti, collected in 1994) were obtained from Prof. A. Failloux via the Infravec2 consortium. For rearing, eggs were hatched in dechlorinated tap water and larvae (±400) were transferred into trays containing 3 L of dechlorinated water and fed daily with a yeast tablet (Gayelord Hauser) until pupation. Pupae were transferred to small bowls and allowed to emerge inside 30 × 30 × 30 cm BugDorm cages (MegaView Science). Adult mosquitoes were supplied with balls soaked in a 10% sucrose solution supplemented with 100 U/mL P/S. The trays and cages were incubated at 28±1°C with a light/dark cycle of 16/8 hours and 80% relative _humidity.104,117,119_

#### 7. Mosquito gut dissection and *ex vivo* culture

Unfed, antibiotic-treated female mosquitoes (3–7 day old) were cold-anaesthetised and surface sterilised by soaking in 70% ethanol for 20 seconds followed by soaking in PBS for 20 seconds. Mosquitoes were dissected in PBS on a petri-dish using the stereomicroscope (VisiScope, VWR), as previously described.^119^ Briefly, the gut was exposed by carefully pulling the second to last segment of the mosquito abdomen. Following gut excision (including the foregut, the midgut, and the hindgut), tracheal tubes and Malpighian tubules were carefully removed. Following dissection, mosquito guts were washed twice in gut medium, consisting of Leibovitz’s L-15 medium supplemented with 2% FBS, 100 U/mL P/S, 50 *µ*g/mL of kanamycin, and 0.25 *µ*g/mL of amphotericin B (Sigma Aldrich), and placed individually in a 96-well tissue culture plate with a clear bottom (PerkinElmer, USA) containing gut medium. Plates were maintained at 28°C, without CO_2_.

#### 8. Viability of *ex vivo* guts following ZP12 treatment

Following gut dissection, the toxicity of ZP12 was assessed by an ATP-based assay using Cell Titer-Glo 3D Reagent, as previously described.^497^ Briefly, dissected mosquito guts were incubated with a range of ZP12 concentrations (10, 50, 75 and 100 *µ*g/mL) or PBS-vehicle, diluted in gut medium for 3 days. Following the ZP12 exposure period, the gut medium was removed and 100 *µ*L of fresh gut medium was added, along with 100 *µ*L of Cell Titer Glo-3D Reagent. The gut-containing plate was agitated on a shaker at 400 RPM for 5 minutes to induce cell lysis. The signal was allowed to stabilise by incubating the plate in the dark, at room temperature for 25 minutes. After incubation, 100 *µ*L of the supernatant was transferred to a black plate and luminescence was recorded using the Tecan plate reader.

#### 9. Infection and antiviral kinetics in mosquito guts

Following mosquito gut dissections, a total of 1×10^5^ PFU/mL of RRV was added to each gut-containing well. Negative controls consisted of dissected guts fixed in 4% PFA for 30 minutes. Guts were incubated for 2 hours at 28°C, without CO_2_. After incubation, the virus inoculum was carefully removed, and guts were washed twice in gut medium. For antiviral studies, ZP12 was diluted in gut medium to either 0.5 µg/mL or 100 µg/mL and added to infected gut cultures. Untreated infected gut cultures served as virus controls. Guts were maintained at 28°C without CO_2_ until collection at the indicated time points.

#### 10. Infectious virion determination

Collected guts were individually homogenised in 250 *µ*L of PBS using bead disruption (Precellys 2.8 mm ceramic beads). The gut homogenate was filtered using 0.8 *µ*m MINI column filters (Sartorius) to remove debris, bacteria and fungi. Infectious virus titres were quantified by TCID_50_ assay. Briefly, Vero cells were seeded at 2.5×10^4^ cells/well in 96-well plates and incubated overnight. Filtered gut homogenates were serially diluted ten-fold and added to Vero cells. Following 72 h incubation at 37°C and 5% CO_2_, wells were scored for cytopathic effect (CPE). TCID_50_/mL values were calculated using the Reed-Muench method.

#### 11. FAIRY: Virucidal mechanism

To assess virucidal activity in infected mosquito tissues, guts were infected and treated as described above. At 3 dpi, guts were removed from culture medium and washed three times in PBS to remove residual extracellular virus. Individual guts were transferred to black 96-well plates containing 50 µL PBS and 50 µL TO-PEG solution. Plates were incubated in the dark for 10 min at room temperature and fluorescence was measured using a Tecan plate reader (*λ*_Ex_=510 nm and *λ*_Em_=533 nm). Multiple fluorescence readings were acquired per well and averaged.

#### 12. Therapeutic treatment of *ex vivo* guts

Mosquito guts were dissected and infected with RRV as described above. To evaluate the therapeutic efficacy of ZP12 against ongoing infection, gut cultures were maintained for 1, 2 or 3 days post-infection (dpi) prior to treatment. At the indicated time point, the culture medium was removed, guts were washed twice with gut medium, and fresh gut medium containing 0.5 µg/mL ZP12 was added. Untreated infected gut cultures received fresh gut medium only. For treatment initiated at 1 dpi, guts were collected after either 24 h (2 dpi) or 48 h (3 dpi) of exposure. For treatment initiated at 2 dpi or 3 dpi, guts were collected 24 h following treatment. At the indicated collection time points, individual guts were harvested, homogenised and infectious viral titres were determined by TCID_50_ assay.

#### 13. Prophylactic treatment of Ae. aegypti mosquitoes

Mosquito rearing was performed as described above, except that adult mosquitoes were supplied with cotton balls soaked in 10% sucrose solution supplemented with 100 U/mL P/S and 100 ng/µL ZP12 for 24 h prior to dissection. Following the feeding period, guts from both ZP12-fed and untreated mosquitoes were dissected and infected *ex vivo* with RRV as described above. To assess whether prior exposure to ZP12 conferred antiviral protection, infectious viral titres were determined at 1, 2 and 3 dpi using TCID_50_ assays.

To determine whether ZP12 remained biologically active within the mosquito following oral administration, whole mosquitoes previously exposed to ZP12 were homogenised in gut medium, filtered, and overlaid onto RRV-infected *ex vivo* gut cultures. Infectious viral titres were quantified at 3 dpi. To investigate whether ZP12 was retained within the crop, crops from ZP12-fed mosquitoes were dissected, homogenised in gut medium, filtered, and overlaid onto RRV-infected *ex vivo* gut cultures. Infectious viral titres were quantified at 3 dpi. To promote mobilisation of stored crop contents, a separate cohort of mosquitoes was starved for 24 h following ZP12 feeding prior to gut dissection and subsequent *ex vivo* RRV infection. Infectious viral titres were quantified at 3 dpi by TCID_50_ assay.

#### 14. Statistical Analysis

For all *in vitro* studies, comparisons between the non-treated (NTC) and treated groups were performed using one-way ANOVA with multiple comparisons. Data were presented as mean±SEM, of three independent experiments performed in at least duplicate, unless otherwise stated. For all *ex vivo* studies, comparisons between experimental groups were performed using a two-tailed non-parametric Mann-Whitney U test. Data are presented as mean±SEM from at least two independent experiments, with 4-6 mosquito guts analysed per group, unless otherwise stated. All statistical analyses were performed using GraphPad Prism (v9.1). Statistical significance was defined as ^∗^p*<*0.05, ^∗∗^p*<*0.01 and ^∗∗∗^p*<*0.001.

