## Supplementary data for "Virucidal Star Polymer Disrupts Alphavirus Infection in Aedes Mosquitoes"

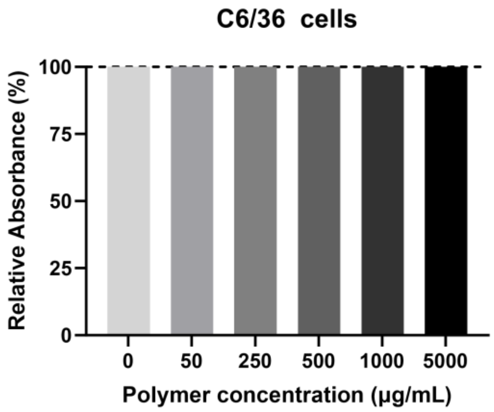


*SI Figure 1: Viability of C6/36 cells assessed using the MTS assay after ZP12 treatment. Percent of viable cells were calculated based on untreated (PBS) cells. Data are presented as means±SEM of three independent experiments performed in triplicate.*
